# Enhancing Functional Connectivity Analysis in Task-Based fMRI Using the BOLD-Filter Method: Greater Network and Activation Voxel Sensitivities

**DOI:** 10.1101/2025.06.18.653042

**Authors:** Yul-Wan Sung, Uk-Su Choi, Motoko Tanabe, Seiji Ogawa

## Abstract

Task-based functional MRI (tb-fMRI) has gained prominence for investigating brain connectivity by engaging specific functional networks during cognitive or behavioral tasks. Compared to resting-state fMRI (rs-fMRI), tb-fMRI provides greater specificity and interpretability, making it a valuable tool for examining task-relevant networks and individual differences in brain function. In this study, we evaluated the utility of the BOLD-filter—a method originally developed to extract reliable BOLD (blood oxygenation level-dependent) components from rs-fMRI—by applying it to tb-fMRI data as a preprocessing step for functional connectivity (FC) analysis. The goal was to enhance the sensitivity and specificity of detecting task-induced functional activity. Compared to the conventional preprocessing method, the BOLD-filter substantially improved the isolation of task-evoked BOLD signals. It identified over eleven times more activation voxels at a high statistical threshold and more than twice as many at a lower threshold. Moreover, FC networks derived from BOLD-filtered signals revealed clearer task-related patterns, including gender-specific differences in brain regions linked to everyday behaviors. These patterns were not detectable using standard preprocessing approaches. Our findings demonstrate that the BOLD-filter enhances the robustness and interpretability of FC analysis in tb-fMRI. By effectively isolating meaningful functional networks, this approach offers significant advantages over conventional preprocessing methods. The BOLD-filter holds promise for advancing both basic neuroscience research and clinical applications by enabling more precise characterization of task-induced brain activity.

## 1. Introduction

Functional magnetic resonance imaging (fMRI) using blood oxygenation level-dependent (BOLD) contrast is the most widely used non-invasive method for assessing human brain function (Ogawa et al., 1990, 1992). Over the past three decades, BOLD fMRI has remained a cornerstone in both basic neuroscience and clinical research, enabling robust and reliable visualization of brain activity.

While resting-state fMRI (rs-fMRI) has been extensively applied to the study of various brain disorders—including dementia, autism spectrum disorder, depression, and schizophrenia (Deng et al., 2024; Giuliani et al., 2024; He et al., 2024; Karim et al., 2024; Li et al., 2013; Müller et al., 2011; Zhao et al., 2024)—its reliance on spontaneous low-frequency signal fluctuations (0.01– 0.1 Hz) poses interpretive challenges. These fluctuations are often assumed to reflect intrinsic neuronal activity (Fox et al., 2006; He et al., 2008), yet they may also contain substantial physiological and non-neuronal noise (Mulpy et al., 2013; Power et al., 2017). To address these limitations, we previously developed the BOLD-filter, a method designed to selectively extract signal components with consistent BOLD-like characteristics, thereby enhancing the interpretability of rs-fMRI connectivity analyses (Sung et al., 2025).

However, functional connectivity (FC) analysis is increasingly being applied to task-based fMRI (tb-fMRI), which offers unique advantages over rs-fMRI. By engaging specific cognitive or behavioral processes, tb-fMRI allows for more targeted exploration of task-relevant neural circuits and provides stronger predictive power for individual differences in performance (Cole et al., 2014; Finn et al., 2015; Gonzalez-Castillo et al., 2018; Greene et al., 2020; Zhao et al., 2023).

In this study, we extend the application of the BOLD-filter to tb-fMRI data to evaluate its effectiveness in enhancing FC analysis during task engagement. Specifically, we assess its performance using multi-echo data and compare the resulting activation maps and region-of-interest (ROI) signals with those derived from a conventional preprocessing pipeline applied to single-echo data (TYP; Chao-Gan & Yu-Feng, 2010). We then construct FC networks from task-responsive ROIs and investigate whether BOLD-filtered data reveal connectivity patterns related to gender differences during a visually guided task with real-world relevance (Zhang et al., 2020; Ingalhalikar et al., 2014). These analyses aim to demonstrate the advantages of BOLD-filter in improving the sensitivity and specificity of task-based FC studies, potentially advancing both research and clinical applications of tb-fMRI.

## 2. Methods

2.0 Methodological framework for investigating the applicability and benefits of BOLD-filter To evaluate the effectiveness of the BOLD-filter—a preprocessing method specifically designed for multi-echo fMRI data—we conducted a comparative analysis of activation maps derived from different preprocessing pipelines. This involved comparing single-echo data processed with a standard preprocessing pipeline (TYP; Chao-Gan & Yu-Feng, 2010) against multi-echo data processed using the BOLD-filter. The goal was to assess the reliability of the BOLD signals obtained under each condition.

In addition to this comparison, we evaluated the performance of the BOLD-filter (Sung et al., 2025) relative to two other established artifact removal techniques for multi-echo data: (1) the simple averaging of multi-echo signals (Posse et al., 1999), and (2) multi-echo independent component analysis (meICA; Kundu et al., 2012). For consistency, each method served as a preprocessing step replacing traditional artifact removal processes, enabling a direct comparison of their ability to improve BOLD signal reliability.

Following the initial evaluation using activation maps, we further assessed the performance of the BOLD-filter in comparison to the TYP pipeline by conducting region-of-interest (ROI) analyses and functional connectivity assessments.

### 2.1 BOLD-filter design

The BOLD-filter method (Sung et al., 2025) was originally developed to identify frequency components exhibiting TE-dependence in resting-state fMRI (rs-fMRI) data, as described in the foundational work on BOLD contrast (Ogawa et al., 1990; 1992). The core idea is to isolate frequency components at each discrete frequency that show a monotonic increase in signal amplitude across multiple echo times (TEs), which serves as an indicator of reliable BOLD signals. Figure 1 illustrates the selection process for TE-dependent frequency components using three echo times.

**Figure 1.**
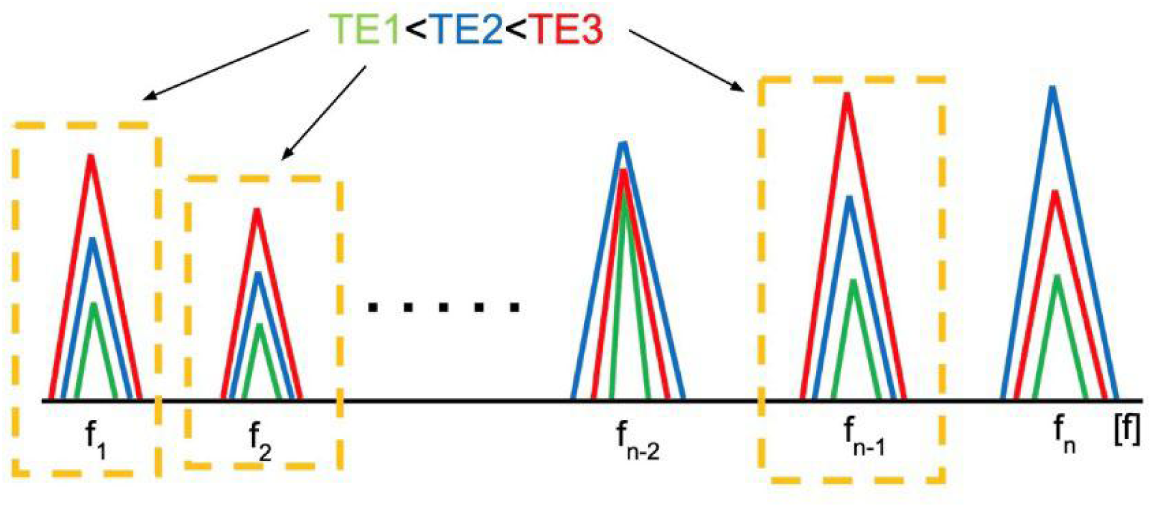
The scheme of the BOLD-filter. The BOLD-filter identifies reliable BOLD components by evaluating the power spectra of three echo signals in the frequency domain, based on the criterion of TE-dependence (i.e., power at TE1 < TE2 < TE3). Here, TE1, TE2, and TE3 correspond to the first, second, and third echo times, respectively. In the plot, green, blue, and red lines represent the power spectra for TE1, TE2, and TE3. Frequencies f1, f2, and ft–1, which satisfy the TE-dependence criterion, are selected and then inverse-transformed to reconstruct the corresponding time-domain signals.

In this study, which utilizes three TEs, we implemented the BOLD-filter by first applying a Fourier transform to the multi-echo fMRI signals. We then applied the following selection criterion in the frequency domain:

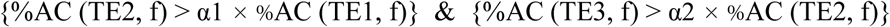

where %AC (TEi, f), i = 1, 2, and3, denotes the percent change of amplitude of signal fluctuations at frequency f for each echo time (TE1, TE2, TE3), and also α1 = 1 + α (TE2/TE1 - 1) and α2 = 1 + α (TE3/TE2 -1) is a threshold parameter that defines the required degree of monotonic increase, serving as a noise rejection margin. In this study, α was set to 0.2, chosen to balance reliability and signal retention for task-based fMRI (tb-fMRI) data.

To further minimize the inclusion of noise-contaminated components, we imposed additional constraints: 1) The Pearson correlation coefficient between the time-domain signals of TE1 and TE2, as well as between TE2 and TE3, was required to exceed r > 0.5, ensuring sufficient coherence across echo times. 2) Conversely, the correlation coefficient between TE1 and TE2, and between TE2 and TE3, was required to remain below r < 0.995 to avoid excessively high coherence that could arise from a dominant noise-driven frequency component or potential artifacts introduced during the Fourier transformation. These correlation checks were performed after applying the inverse Fourier transform to the selected frequency components. In addition, a high-pass filter with a cutoff frequency of 0.008 Hz was applied to further reduce potential artifacts introduced during the Fourier transformation.

Unlike in our prior work on rs-fMRI, we did not impose an additional criterion based on the variation of time courses. In the context of tb-fMRI, where a stimulus pattern is available as a reference (i.e., ground truth), this constraint was deemed unnecessarily strict and was omitted to preserve task-related BOLD signal components.

The MATLAB implementation of the BOLD-filter, including additional minor details, is provided in the Supplementary Methods.

### 2.2 MRI Measurements

All MRI experiments were conducted using a Skyra-fit 3 Tesla MRI scanner (Siemens, Germany) equipped with a standard 20-channel head matrix coil.

#### (1) fMRI Imaging

##### - MRI Parameters

A multi-echo echo planar imaging (EPI) sequence was used for functional MRI, modified from the default single-shot, three-echo gradient-echo EPI (GE-EPI) sequence provided by Siemens. This modification enabled acquisition at three echo times: 9.98 ms, 21.61 ms, and 33.24 ms. The maximum TE of 33.24 ms was selected to capture monotonically increasing BOLD signals. The multi-echo GE-EPI sequence had the following acquisition parameters: Repetition time (TR): 1000 ms, Flip angle: 70 degrees, Field of view (FOV): 220 mm, Matrix size: 64 × 64, Slice thickness: 3.4 mm, with no gap, Multiband acceleration factor: 2, Number of slices: 34, acquired parallel to the anterior commissure–posterior commissure (AC-PC) line. The total scan duration was 243 seconds, including an initial 33-second rest period followed by six alternating blocks of 15-second stimulation and 20-second rest periods.

##### - Task Presentation

During each 15-second stimulation period, movie clips depicting daily behaviors were presented.

These included scenes such as lifting and relocating a heavy object, walking up stairs, vacuuming the floor, and walking down a hallway. During the rest periods, a grey cross-hair was shown at the center of a black screen (Supplementary Fig. 1). The visual stimuli were presented with a visual angle of approximately 30 × 20 degrees.

#### (2) Structural Imaging

Following functional imaging, T1-weighted anatomical images were acquired using an inversion recovery and magnetization-prepared rapid acquisition with gradient echo (MPRAGE) sequence. Imaging parameters included a matrix size of 256 × 256, a field of view (FOV) of 256 mm, and a slice thickness of 1 mm.

### 2.3 Subjects

For the main resting-state fMRI (rs-fMRI) experiments, twenty healthy volunteers participated (mean age: 20.9 ± 0.9 years; 10 males, 10 females). There was no significant age difference between males and females (p = 0.6). Participant health was confirmed by physical examinations conducted within six months prior to the experiments. All participants had no history of neurological disorders or medical conditions contraindicating MRI (e.g., pregnancy, pacemakers, or claustrophobia), as determined by pre-screening interviews. Written informed consent was obtained from all participants in accordance with the Declaration of Helsinki. The study protocol was approved by the Institutional Review Board.

### 2.4 Data Analysis

Activation maps, time courses, and functional connectivity were analyzed from task-based fMRI (tb-fMRI) data following preprocessing, using both multi-echo time series (from all three echo times) and single-echo time series corresponding to the third echo time (TE3).

#### (1) Preprocessing

-BOLDFLT: tb-fMRI data were preprocessed using the BOLD-filter method, which included the following steps: 3D motion correction, BOLD-filter processing, co-registration with individual structural images, normalization, and spatial smoothing using a 4-mm full width at half maximum (FWHM) Gaussian filter. The BOLD-filter algorithm was implemented in MATLAB (MathWorks Inc., Natick, MA, USA) and the other processing was performed using the Data Processing Assistant for Resting-State fMRI (DPABI) software (Chao-Gan & Yu-Feng, 2010; Yan et al., 2016).

- TYP: The standard preprocessing pipeline included 3D motion correction, regression of head motion effects using the Friston 24-parameter model, band-pass filtering (0.01–0.1 Hz) that preserves task-induced signals in block designs such as in this study, and artifact removal based on cerebrospinal fluid (CSF) signals. Images were then spatially smoothed with a 4-mm FWHM filter and co-registered to corresponding anatomical images. All preprocessing was performed using the Data Processing Assistant for Resting-State fMRI (DPABI) software (Chao-Gan & Yu- Feng, 2010; Yan et al., 2016).

- Both BOLDFLT and TYP methods shared common steps such as motion correction, normalization, and spatial smoothing.

#### (2) Activation Maps

A general linear model (GLM) was applied to the time series of each voxel, using a two-gamma hemodynamic response function (HRF) as the regressor. This analysis was performed independently for each subject. To visualize voxels showing consistent activation across subjects, spatial probability maps were generated using Cohen’s d after calculating the distance for each subject (Park et al., 2004). Separate maps were constructed for both BOLDFLT and TYP data.

#### (3) ROI-Based Analysis

A brain atlas comprising 48 Brodmann areas (BA) was used for region-of-interest (ROI) analysis due to the well-established functional roles of these areas. For each ROI, the time course was averaged and correlated with the reference (task) regressor using Pearson’s correlation. ROIs showing a correlation coefficient greater than r = 0.5 were considered to exhibit task-induced activity and were selected for subsequent functional connectivity analysis.

#### (4) Functional Connectivity and Gender Association

Functional connectivity was assessed using Pearson’s correlation between ROI signals among ROIs identified as task-induced. Edges of the functional connectivity network were defined by statistically significant correlations (p < 0.05). Associations between these edges and gender were evaluated using Spearman’s correlation.

#### (5) Supplementary analysis: activation maps

##### 1) Averaging of multi echo Time Series (AVG)

To create an averaged time series, the signals from TE1, TE2, and TE3 were first normalized by the mean value of the TE3 time series. These normalized signals were then averaged to form a composite time series. This averaged time series underwent the same preprocessing steps as those used in the TYP pipeline. Subsequently, GLM was applied to the averaged time series for each voxel, utilizing a two-gamma hemodynamic response function (HRF) as the regressor. This analysis was conducted independently for each subject.

##### 2) Multi-Echo Independent Component Analysis (meICA)

The meICA method (Kundu et al., 2012), a well-established artifact removal technique for multi-echo fMRI data, was applied to the TE1, TE2, and TE3 time series. This method is implemented within the *tedana* software package (https://tedana.readthedocs.io/en/stable/approach.html). The resulting denoised time series were then subjected to normalization and spatial smoothing procedures, consistent with those used in the BOLDFLT pipeline.

## 3. Results

The activation maps generated by stimulation revealed that BOLDFLT produced significantly more statistically reliable voxels compared to TYP. Figure 2 presents activation maps from a representative subject, showing a substantial overlap between BOLDFLT and TYP at a low statistical threshold (t-value = 4). However, at a higher threshold (t-value = 10), BOLDFLT activation extended across a broader range of brain regions than TYP (Fig. 3). This trend varied between subjects. For example, in another representative subject, BOLDFLT detected numerous activation voxels at the high threshold, while TYP identified none (Supplementary Fig. 2 and 3). This subject exhibited an even greater difference between BOLDFLT and TYP than the first.

**Figure 2.**
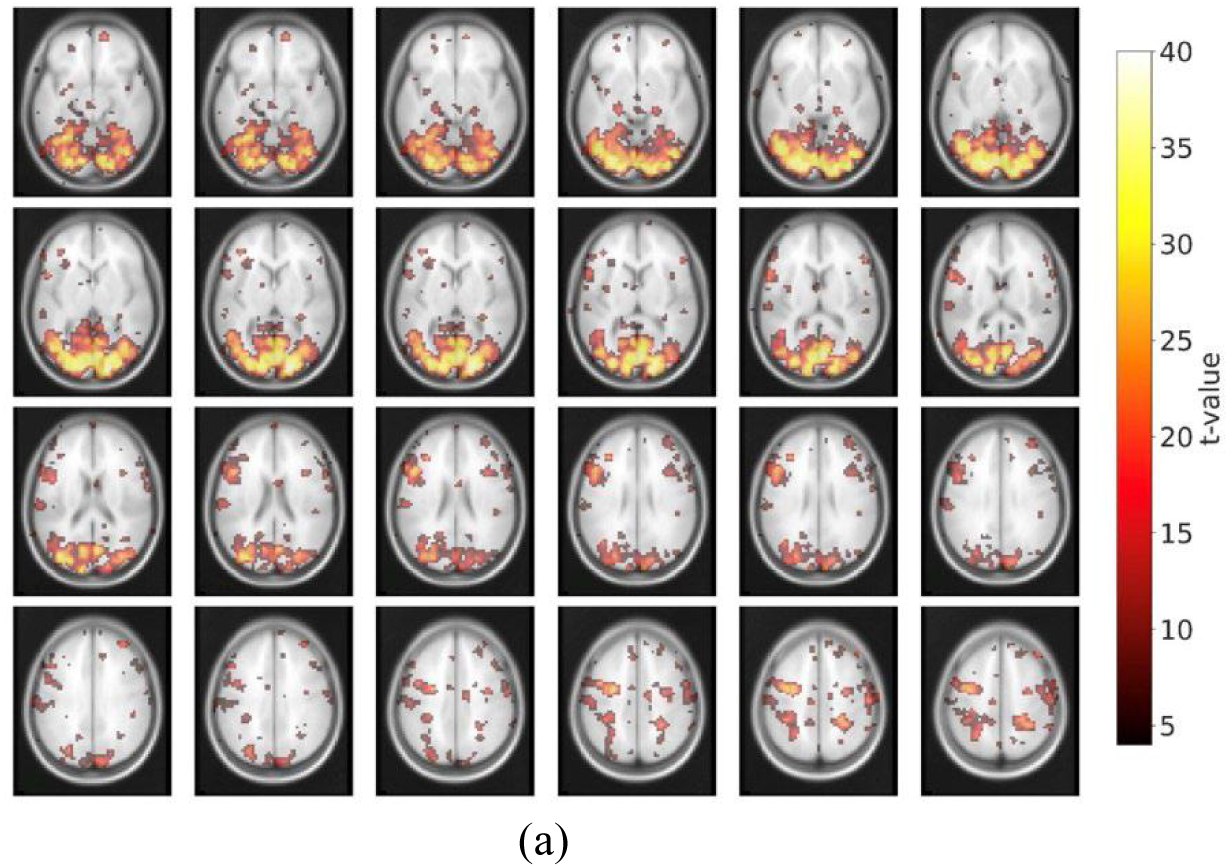
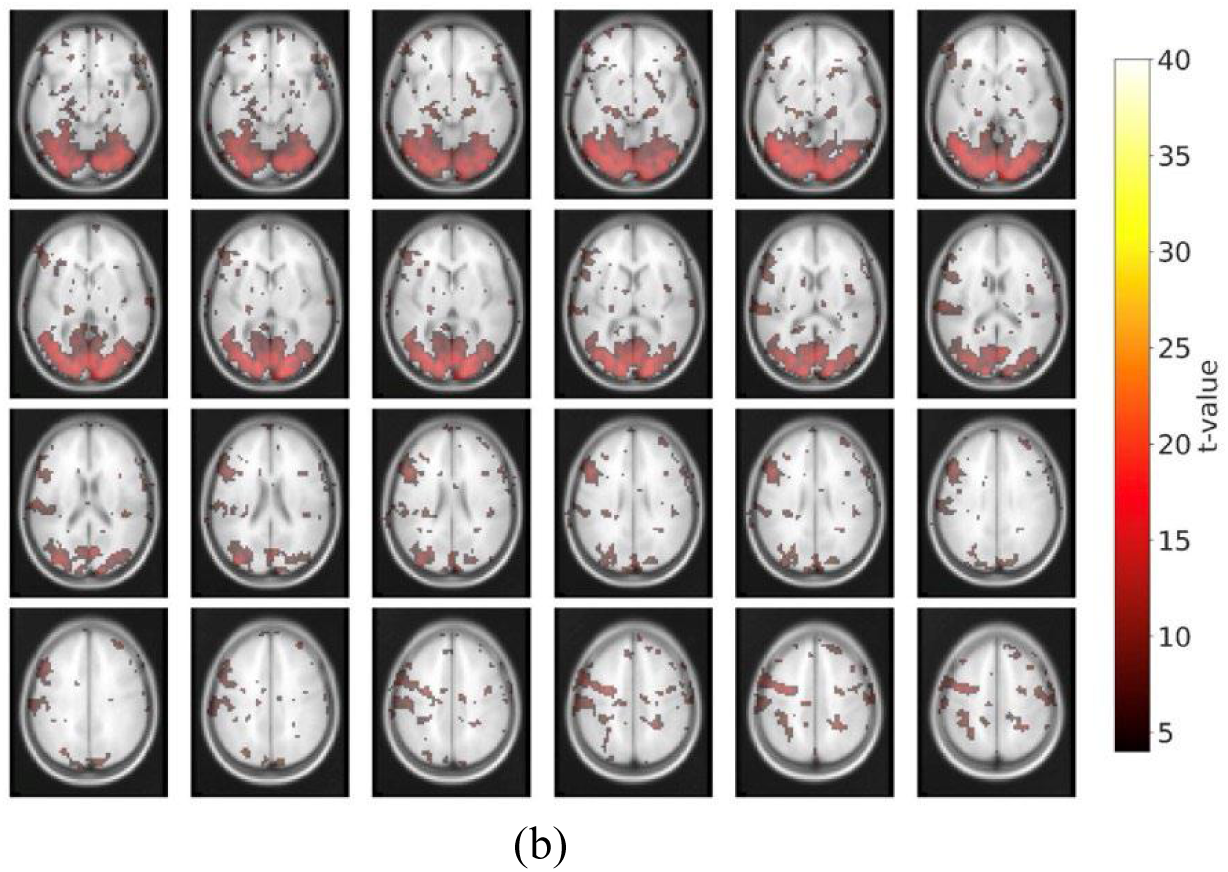
Activation maps. This figure presents activation maps from a representative subject at a low threshold (t = 4). The color bar indicates *t*-values. (a) Activation map obtained using BOLD-filter preprocessing (BOLDFLT); (b) Activation map obtained using conventional preprocessing (TYP).

**Figure 3.**
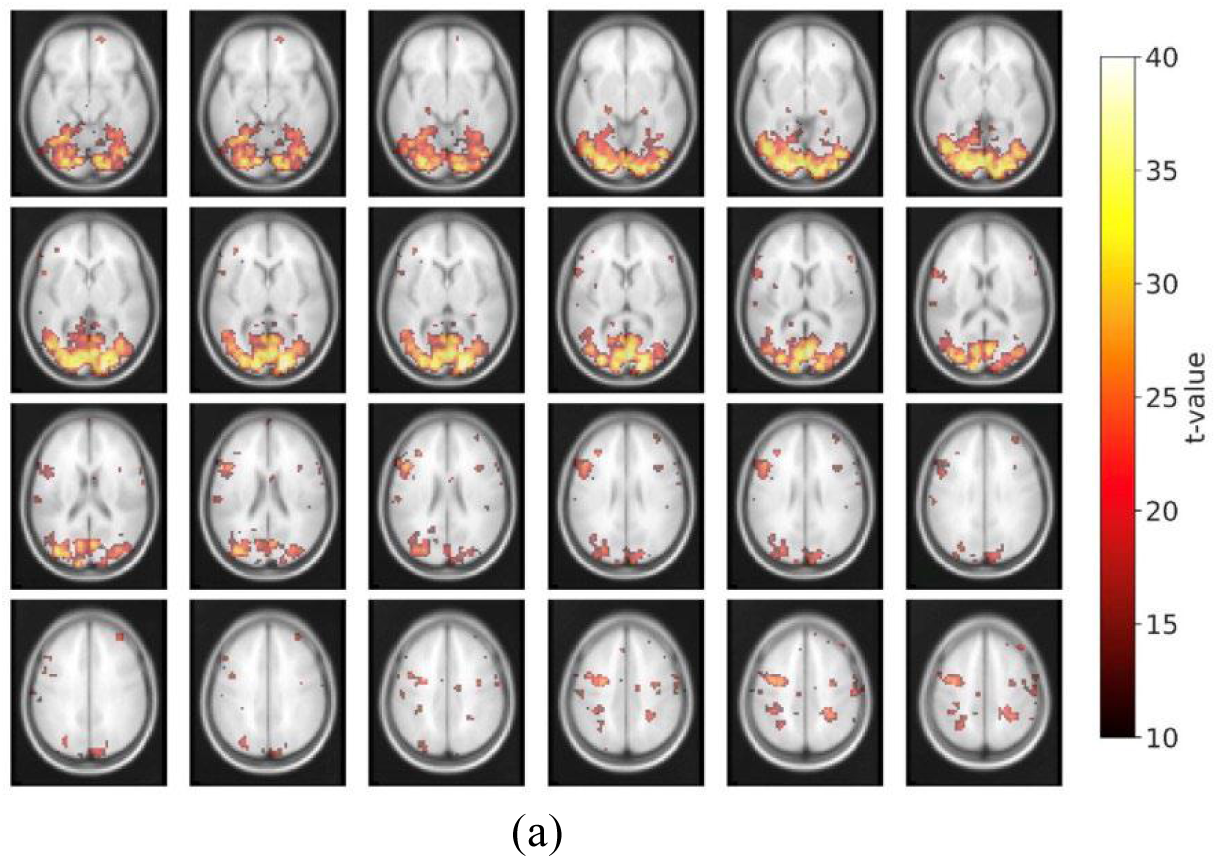
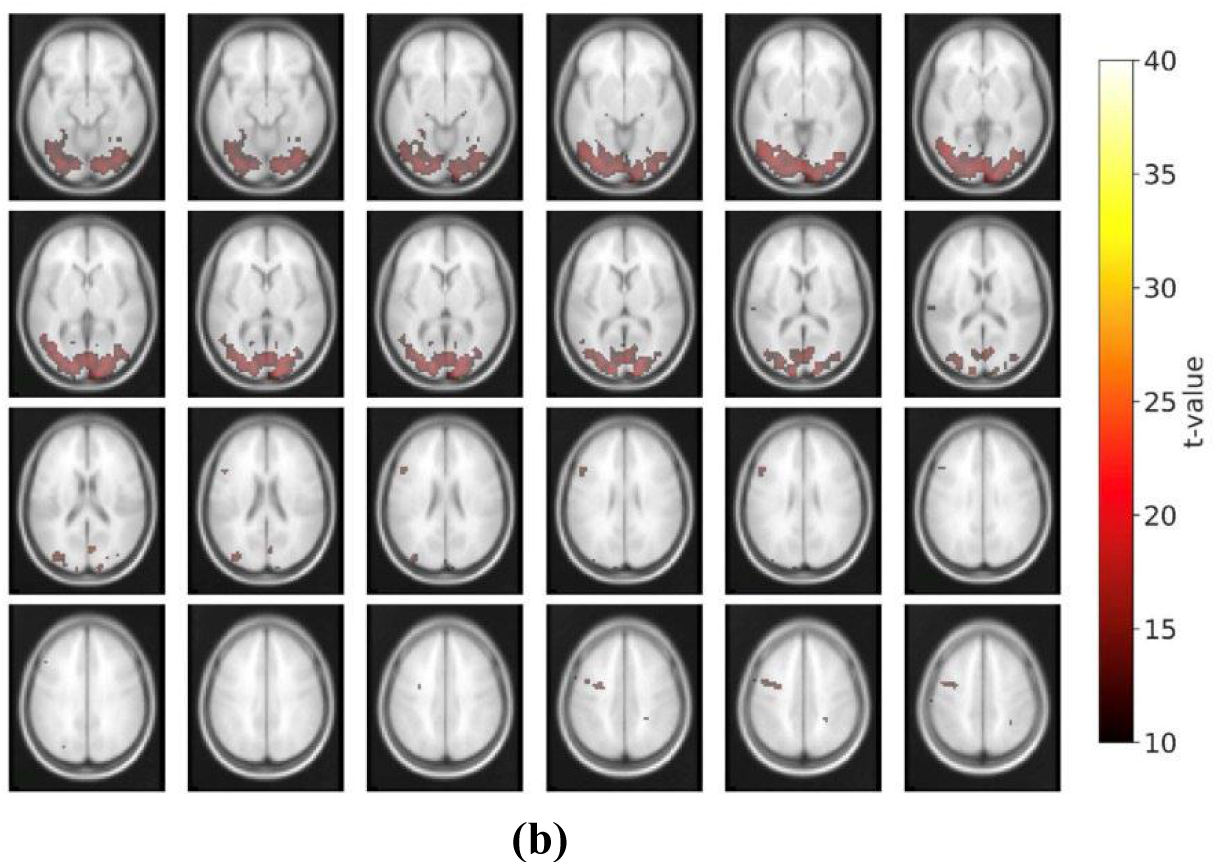
Activation maps. This figure presents activation maps from the representative subject at a high threshold (t = 10). The color bar indicates *t*-values. (a) Activation map obtained using BOLD-filter preprocessing (BOLDFLT); (b) Activation map obtained using conventional preprocessing (TYP).

To quantify these activation profiles across all subjects, we calculated Cohen’s d (Swinton et al., 2023) to represent activation intensity and created corresponding probability maps. These maps demonstrated that activation intensity for BOLDFLT was consistently higher than for TYP and mirrored the patterns observed in individual activation maps. Notably, BOLDFLT revealed stronger activation beyond the primary visual regions (Fig. 4).

**Figure 4.**
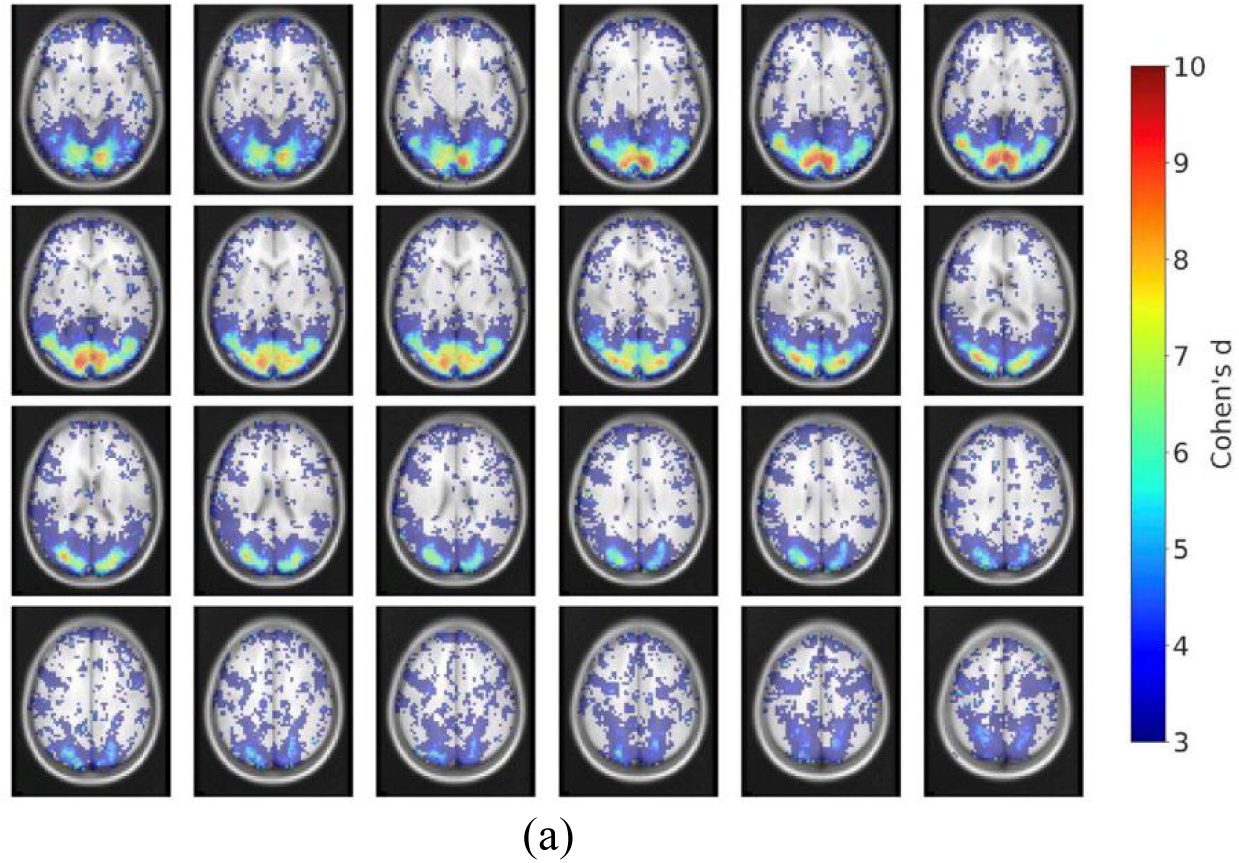
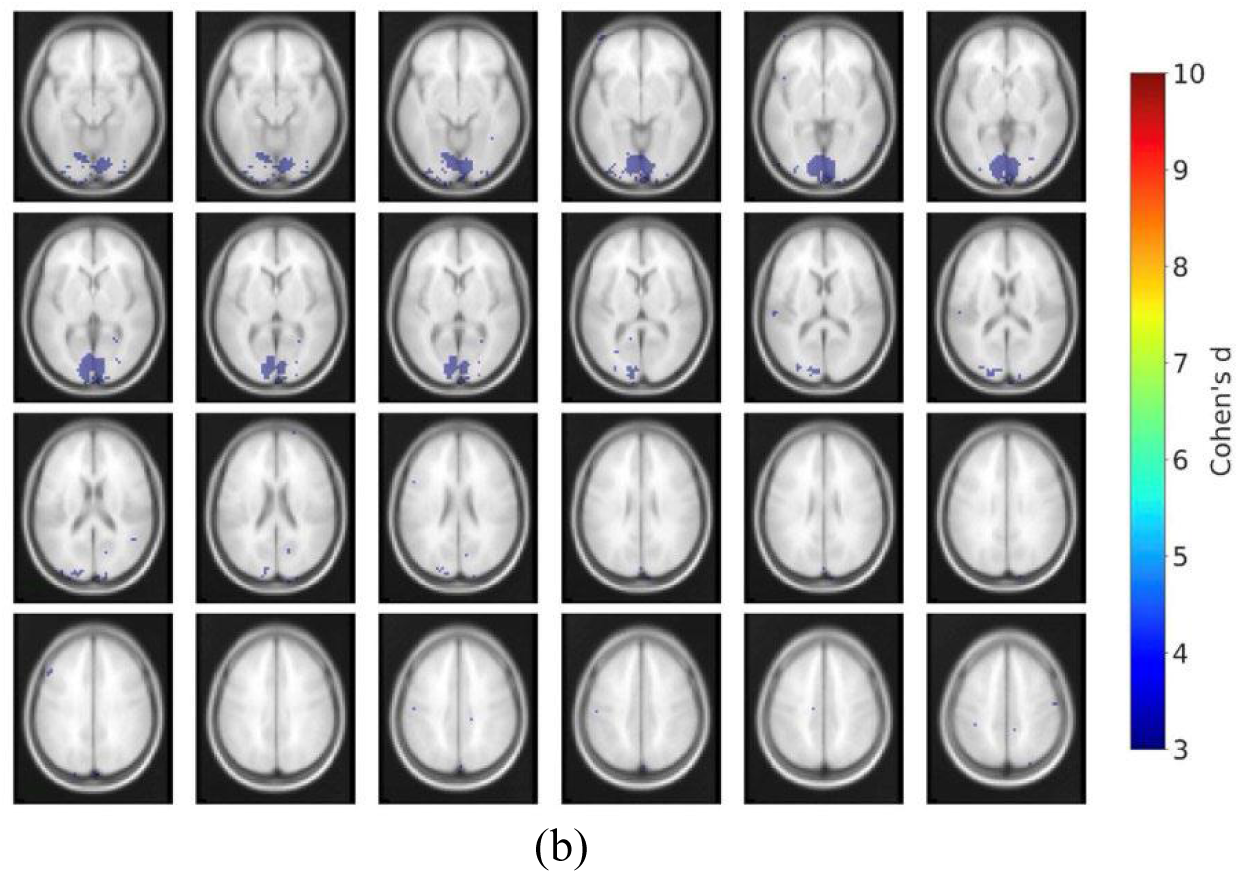
Group-level probability maps. Probability maps generated from activation data across all subjects, thresholded at Cohen’s *d* = 3. (a) BOLD-filter preprocessing (BOLDFLT); (b) Conventional preprocessing (TYP). The color bar indicates Cohen’s *d* -values.

BOLDFLT produced significantly more activation voxels than TYP at both high (t=10) and low (t=4) statistical thresholds (paired t-test: *p* = 3.53×10^−8^ and *p* = 3.25×10^−5^, respectively; Fig. 5). Specifically, at the high threshold (t=10), the mean number of voxels (± SEM) was 9,198.9 ± 854.2 for BOLDFLT versus 782.1 ± 223.7 for TYP. At the low threshold (t=4), the means were 17,153.0 ± 1,212.3 for BOLDFLT and 8,283.0 ± 948.0 for TYP.

**Figure 5.**
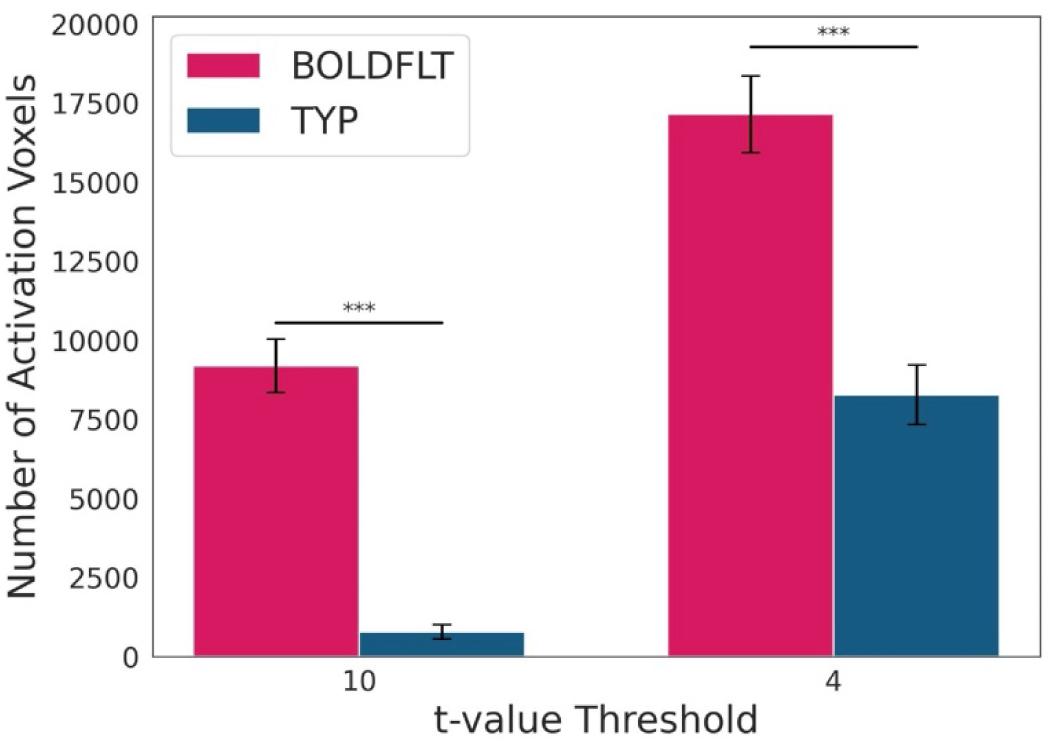
Number of activation voxels. Activation voxels were quantified for BOLD-filter preprocessing (BOLDFLT) and conventional preprocessing (TYP) at both high and low statistical thresholds. At the high threshold (t = 10), BOLDFLT yielded more than 11 times the number of activation voxels compared to TYP. At the low threshold (t = 4), the voxel count for BOLDFLT was more than double that of TYP. (a) BOLDFLT; (b) TYP.

A comparison was also made with the AVG method, which approximates a weighted summation of signals from the three echo times, processed using the same preprocessing steps as TYP. BOLDFLT still yielded significantly more activation voxels than AVG at both high (t=10; *p* = 1.74×10^−7^) and low (t=4; *p* = 0.020) thresholds (paired t-test; Supplementary Fig. 4). The mean number of activation voxels (± SEM) detected by AVG was 3,478.0 ± 350.3 at t=10 and 13,119.0 ± 1078.9 at t=4.

An additional comparison was conducted using the meICA method, a denoising approach that utilizes signals acquired at three different echo times. Even in this comparison, BOLDFLT produced significantly more activation voxels than meICA at both high (t = 10; *p* = 580 × 10^−7^) and low (t = 4; *p* = 0.002) statistical thresholds based on paired *t*-tests (Supplementary Fig. 5). The mean number of activation voxels (± SEM) detected by meICA was 2,977.4 ± 654.9 at t = 10 and 10,596.0 ± 1929.5 at t=4.

To prepare for functional connectivity analysis, we first examined task dependence of signals in the 41 Brodmann areas (ROIs). For each ROI, the average signal across all subjects was computed and then correlated with the reference time course (i.e., the regressor for task stimulation pattern) (Supplementary Fig. 6). We used a correlation threshold of *r* = 0.5 to classify signals as task-induced.

Out of the 41 ROIs, 28 showed correlation values greater than 0.5 for BOLDFLT (red plots above the dotted line in Fig. 6), whereas only 8 ROIs exceeded this threshold for TYP (green plots above the dotted line in Fig. 6). These results suggest that BOLDFLT was more effective at capturing task-induced signal changes across a broader range of brain regions.

**Figure 6.**
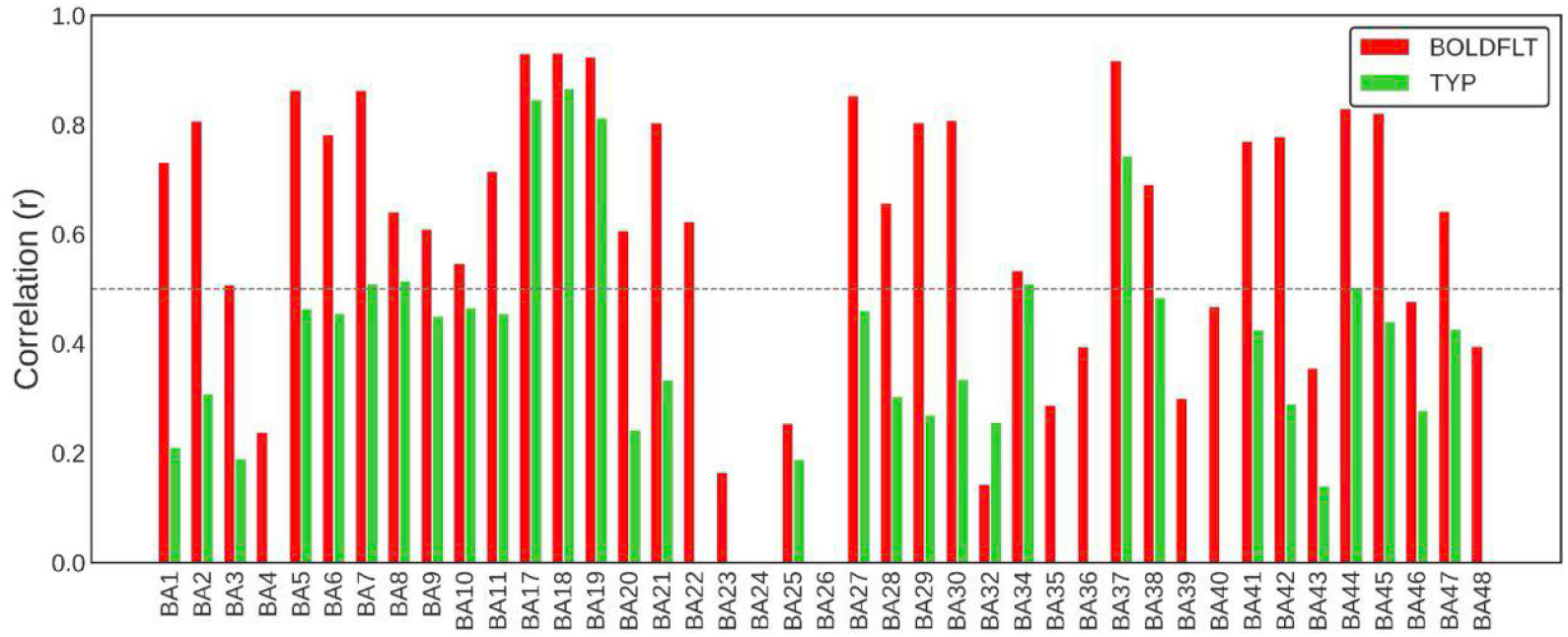
Correlation values at ROIs. Correlation values between the reference time course and each Brodmann area are shown for BOLD-filter preprocessing (BOLDFLT) and conventional preprocessing (TYP). Only regions with non-significant differences (p > 0.05) are included. Red bars indicate BOLDFLT, and green bars indicate TYP.

The ROI time courses across nearly all Brodmann areas—except one—more closely followed the reference time course under BOLDFLT than under TYP, as evidenced by higher correlation values (Fig. 6). Representative time courses from both low- and high-order cortical regions are presented in Figure 7. In the primary visual areas (BA 17, 18, and 19), BOLDFLT and TYP showed similar signal patterns (Fig. 7a). However, in higher-order frontal areas (BA 44, 45, and 46), BOLDFLT time courses closely followed the reference time course, while those of TYP did not (Fig. 7b).

**Figure 7.**
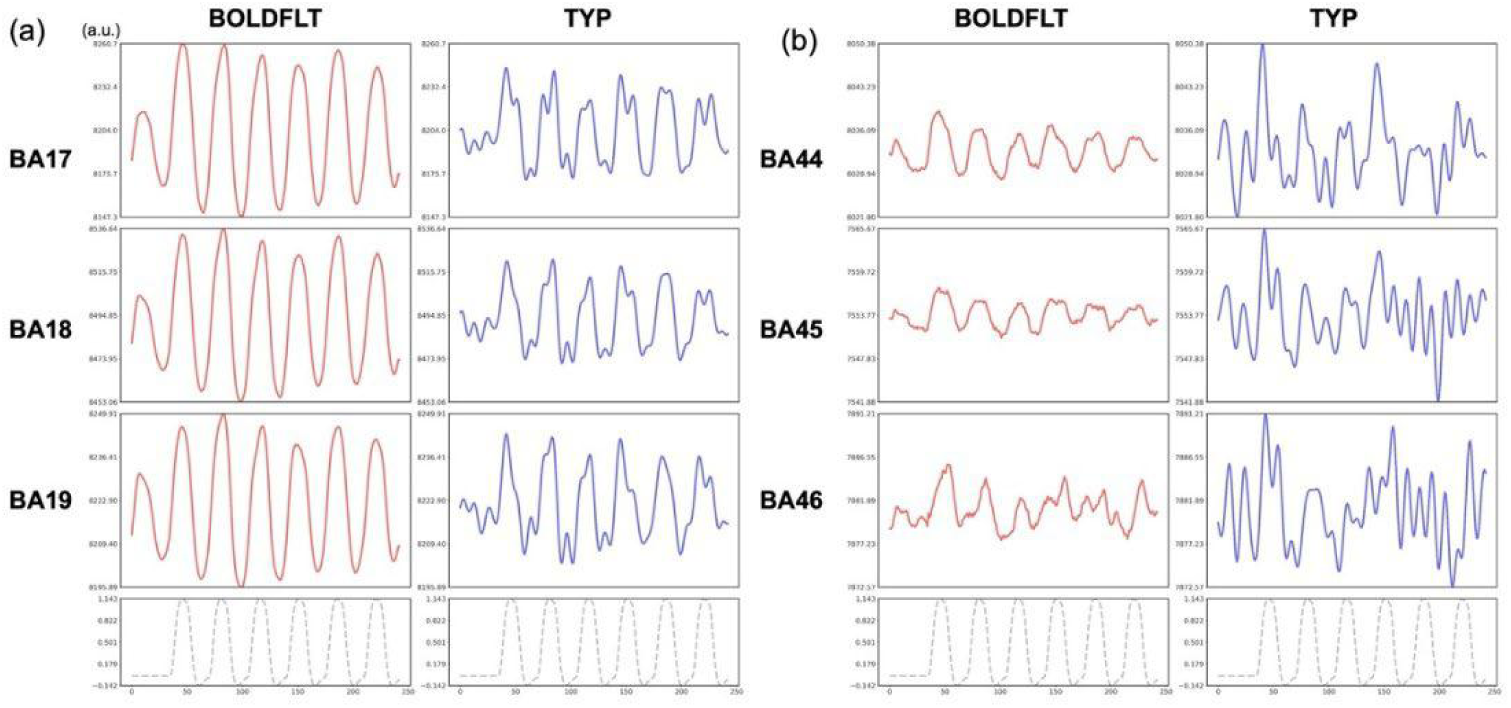
Time course signals. Time course signals averaged across all subjects at representative ROIs for BOLDFLT and TYP. Blue color plots represent TYP and red color plots represent BOLDFLT. Dashed line plots represent the reference time course (task regressor). (a) primary and secondary visual areas: BA17, 18 and 19, (b) high order areas at the frontal region: BA44, 45, 46.

The amplitude differences between BOLDFLT and TYP in these higher-order areas further highlight BOLDFLT’s ability to isolate task-induced BOLD components from mixed signals that also include task-independent fluctuations (Fig. 7b). This contrast was even more apparent in individual subject data (Supplementary Figs. 7 and 8).

Given that BOLDFLT identified more task-relevant signals than TYP, we proceeded to analyze task-induced functional connectivity. Functional connectivity maps were constructed using only the ROIs with *r* > 0.5 correlations to the reference time course (Fig. 8). Both BOLDFLT (Fig. 8a) and TYP (Fig. 8b) networks showed dense interconnectivity among these ROIs, likely due to shared stimulation-related signal changes. However, BOLDFLT demonstrated stronger connectivity values even between several common regions, including BA 17, 18, 19, and 37.

**Figure 8.**
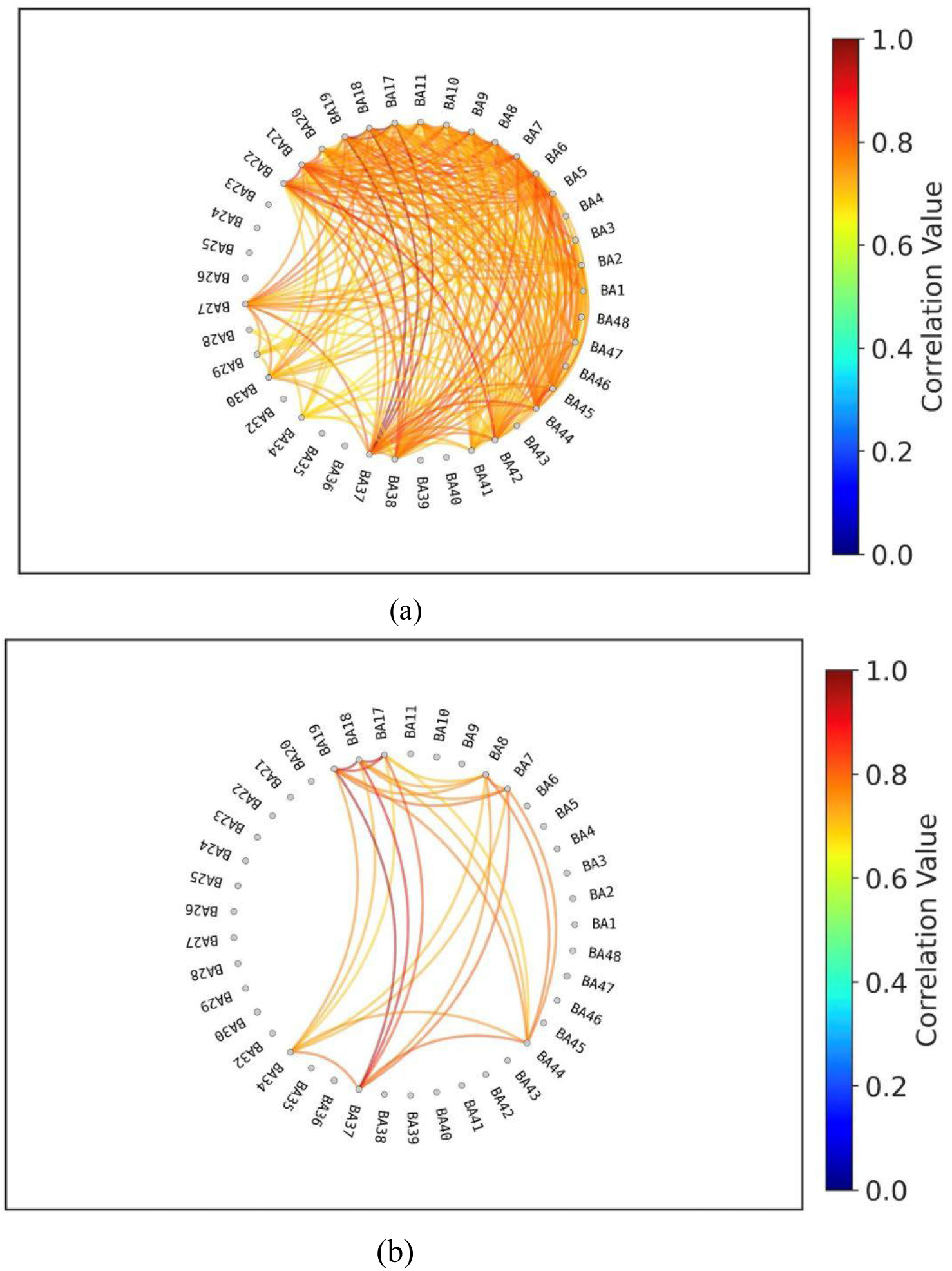
Functional connectivity maps. Functional connectivity maps were constructed using only regions of interest (ROIs) that showed a correlation coefficient greater than *r* > 0.5 with the reference time course. The color bar indicates *r*-values. (a) BOLD-filter preprocessing (BOLDFLT); (b) Conventional preprocessing (TYP).

Analysis of the association between task-induced functional connectivity and gender, with data processed using TYP, revealed no significant connections. In contrast, processing with BOLDFLT identified gender-related connectivity differences in 9 regions of interest (ROIs). (Fig. 9). These significant connections included: <BA 1 – BA 27, 41>, <BA 7 – BA 46>, <BA 9 – BA 37, 44>, <BA 10 – BA 18, 19, 37>.

**Figure 9.**
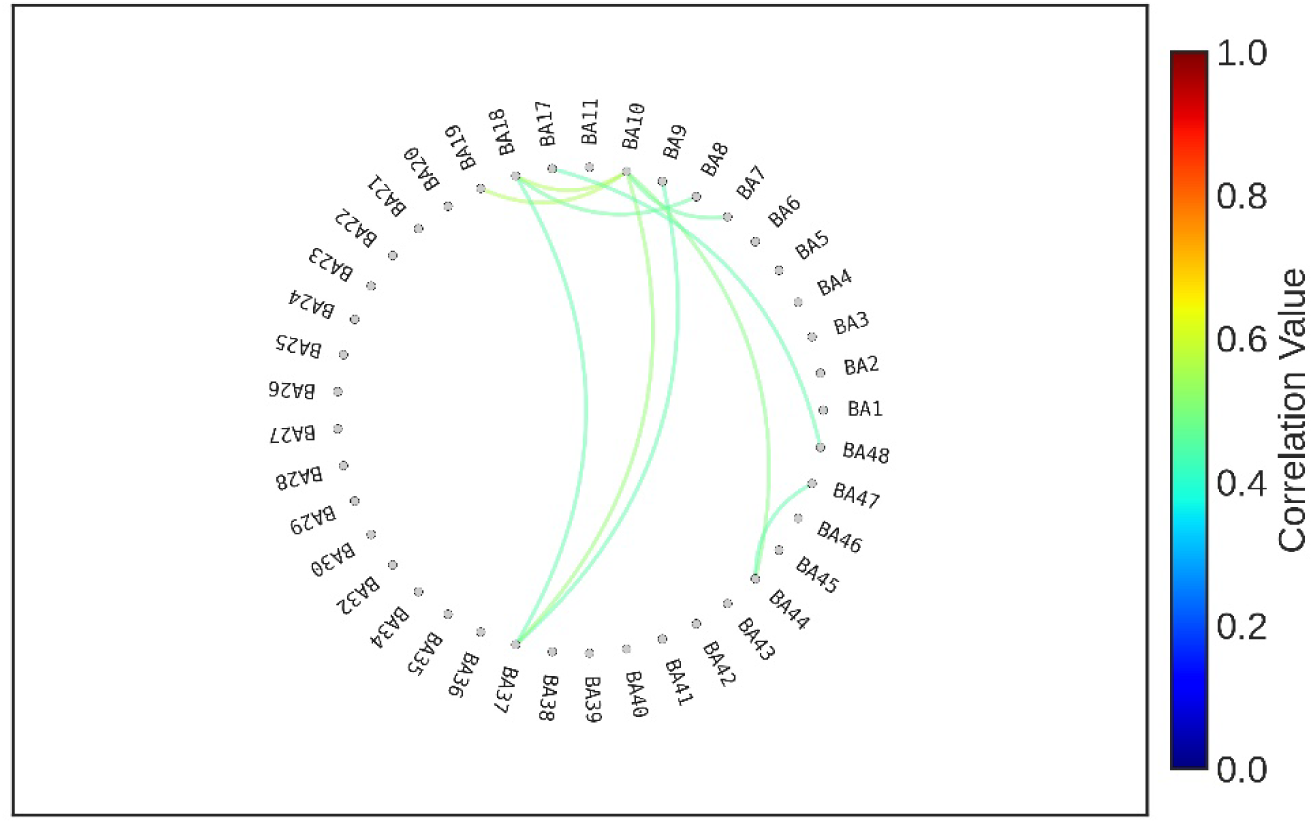
Gender association of functional connectivity. The network was constructed by identifying gender-associated edges within the functional connectivity network. This connectivity network was based on regions of interest (ROIs) that exhibited correlation coefficients greater than *r* > 0.5 with the reference time course.

These results indicate that gender differences in task-induced functional connectivity involve both frontal regions (BA 9, 10, 44, 47, 48) and visual areas (BA 18, 19, 37). Among these, BA 10 and BA 37 emerged as functional connectivity hubs, each forming three or more significant gender-related connections.

## 4. Discussion

In this study, we applied the BOLD-filter to task-based fMRI (tb-fMRI) data to evaluate its effectiveness in extracting reliable BOLD signals. The results validated the superior performance of BOLDFLT by identifying a substantially larger number of task-induced activation voxels and significantly task-modulated signals across a wider range of regions of interest (ROIs), compared to TYP, AVG and meICA. Furthermore, in case of BOLDFLT, functional connectivity analyses based on the ROIs with verified task-induced signals revealed gender-related differences in higher-order brain networks involved in processing the visual tasks, highlighting the potential of BOLD-filter for uncovering meaningful brain-behavior associations.

The number of activation voxels elicited by the visual stimulation was notably higher for BOLDFLT compared to TYP, as well as relative to AVG and meICA across both liberal and conservative statistical thresholds. At the more stringent threshold, the number of activation voxels identified by BOLDFLT was more than 11 times higher than that of TYP, and over twice as high at the lower threshold (Fig. 5) (Dowdle et al., 2023). These results demonstrate the BOLD- filter’s enhanced sensitivity in extracting BOLD components associated with task-induced neural activity. Further, among 41 analyzed Brodmann area (BA) ROIs, the ROI time courses from 28 ROIs showed significant task-related modulation following BOLDFLT processing, whereas only 8 ROIs reached significance after TYP preprocessing (Fig. 6) (Gerchen & Kirsch, 2017). This again highlights the superior ability of the BOLD-filter to retain meaningful BOLD signals from fMRI time courses contaminated with noise.

Functional connectivity computed from these task-relevant ROIs reflects task-driven co-activation across brain regions. Since these ROIs were selected based on their significant responses to task stimulation, the resulting connectivity maps are more likely to represent functionally meaningful, stimulus-driven interactions (Cole et al., 2014). This led to a denser and more coherent functional network under BOLDFLT compared to TYP, consistent with the greater number of ROIs with reliable task-induced signals.

Interestingly, while TYP did not identify any significant connections related to gender differences—resulting in no observable functional network—, BOLDFLT revealed a functional network involving 11 ROIs associated with gender (Satterthwaite et al., 2014). The most prominent hubs within this network were BA 10 and BA 37. BA 37, located in the occipitotemporal cortex, encompasses the fusiform face area and fusiform body area—regions known for processing faces and biological motion. BA 10, part of the anterior prefrontal cortex, is implicated in complex cognition and multitasking (Kanwisher et al., 1997; Koechlin et al., 1999; Peelen & Downing, 2005). Additional gender-sensitive connections included: BA 8 with BA 18; BA 44 with BA 47. These connections collectively represent brain systems involved in interpreting biological motion, emotional visual processing, executive control, spatial localization, motor coordination, mirroring other’s emotion/behaviors and memory integration (Iacoboni &

Mazziotta, 2007; Aminoff et al., 2013; Ochsner et al., 2002). The functional roles of these areas align well with the behavioral content in the visual stimuli, such as "lifting and relocating a heavy object," "going up stairs," "vacuuming the floor," and "walking down a hallway," all of which involved dynamic human body movement and faces (Iacoboni & Mazziotta, 2007; Kanwisher et al., 1997; Satterthwaite et al., 2014). The observed gender differences in these networks may reflect differential cognitive and emotional processing of the same task stimuli. One possible explanation is that physically demanding tasks, such as lifting heavy objects, may be experienced as more challenging by females, while tasks like vacuuming might be more familiar, influencing brain responses and connectivity patterns accordingly (Zhang et al., 2020; Ingalhalikar et al., 2014).

Signal fluctuations observed prior to stimulation onset—above the baseline—may reflect the initial effects of MRI scanning, such as acoustic noise from the scanner, or artifacts introduced during the Fourier transform process (Tomasi et al., 2005). However, the similarity of these pre-stimulus fluctuations in both preprocessing pipelines, particularly their prominence in individual subjects under the TYP method (as shown in Supplementary Fig. 5 and 6), suggests they are more likely attributable to scanner-induced noise rather than artifacts from frequency transformation. Although the impact of these initial fluctuations on the current results is likely negligible, removing the initial period—approximately 10 seconds—before conducting functional connectivity analysis, as is commonly done in rs-fMRI studies, may also be a beneficial approach in task-based fMRI (Soares et al., 2016).

The higher signal amplitudes observed in low-order visual areas (BA 17, 18, and 19) after BOLDFLT processing, compared to TYP, may be due to more faithful preservation of the stimulus-induced signal shape (Fig. 7a). In BOLDFLT, the signal changes closely track the reference time course, whereas in TYP, the same signals appear to be degraded or attenuated by noise fluctuations. This difference is likely amplified by the averaging process across subjects and ROIs.

Ideally, signals extracted by BOLD-filter should closely follow the reference time course, resulting in minimal differences in activation maps between low and high statistical thresholds. However, even with BOLDFLT, the number of activated voxels at the high threshold was approximately 1.8 times lower than at the low threshold. This indicates that BOLD-filter, while effective, is still susceptible to a degree of noise. One way to reduce this gap is by increasing the filter parameter α beyond the value of 0.2 used in this study. However, a higher α value would impose stricter criteria for BOLD component selection, potentially reducing the number of detected activation voxels—a trade-off between signal reliability and sensitivity.

In task-based fMRI, using a lower α value can be justified because the reference time course (i.e., the known stimulation pattern) allows for additional filtering, such as computing correlations between ROI signals and the reference. Despite the possibility of residual noise even after BOLD-filtering, our findings demonstrate that BOLD-filter consistently outperforms the conventional TYP preprocessing (Chao-Gan & Yu-Feng, 2010; Yan et al., 2016). This supports its utility for enhancing signal quality and improving the reliability of functional connectivity analysis in task-based fMRI studies.

Several multi-echo analysis methods have been proposed in previous studies (Posse et al., 1999; Poser et al., 2006; Kundu et al., 2012). One straightforward approach involves using an average of signals from the three echo times or fitting them using a model-based technique (Posse et al., 1999), both of which have been shown to enhance sensitivity. Another widely used method is multi-echo independent component analysis (meICA) (Kundu et al., 2012), which aims to separate and remove non-BOLD components based on their TE-independence. In contrast, BOLD-filter is designed to extract BOLD components that meet specific TE-dependence criteria, focusing on enhancing the reliability of BOLD signals rather than removing noise per se. While both meICA and BOLD-filter contribute to improving the signal-to-noise ratio, their underlying goals and mechanisms differ. In a previous study, we demonstrated that the BOLD-filter outperformed meICA in identifying reliable BOLD components, based on a framework that defines BOLD reliability through TE-dependence and temporal coherence across the three echo time series (Sung et al., 2025). In the current study using task-based fMRI data, the BOLD-filter again substantially outperformed both meICA and the averaging method in detecting task-induced BOLD signals, as evidenced by the number of activation voxels identified (Supplementary Figs. 4 and 5). These results provide further support for the BOLD-filter’s effectiveness in extracting robust task-related functional signals.

The Brodmann’s areas used as ROIs clearly reflected task-induced signals in the lower-order visual regions—specifically BA 17, BA 18, and BA 19—which are well-established as primary and secondary visual cortices (Wandell et al., 2007). This validates the use of Brodmann’s areas as a suitable atlas for our analysis. However, employing a more finely parcellated brain atlas could allow for a more detailed examination of functional connectivity networks, potentially capturing more nuanced region-specific interactions (Lopez-Lopez, et al., 2020).

Various preprocessing pipelines exist, and applying them after BOLD-filtering may further enhance performance. Therefore, we compared BOLD-filter with TYP, which includes the most commonly used preprocessing steps. This comparison supports our rationale for using TYP as a reference, as adding additional preprocessing steps to TYP would likely also improve performance for single-echo signals.

This study has several limitations that should be acknowledged. First, the task involved passive viewing of video clips, which limits the ability to pinpoint which specific behaviors contributed to the observed gender differences in functional connectivity (Satterthwaite et al., 2014). Second, the sample size may have been insufficient to capture the full range of individual variability or to ensure the generalizability of the gender-related findings in the functional connectivity networks. Third, the scan duration might have been relatively short for robust functional connectivity analysis, as longer scans are typically recommended (Birn et al. 2013). However, extending scan durations during task performance may introduce increased physiological variability, which could confound the results (Krüger & Glover, 2001). Fourth, some parameters used in the BOLD-filter were determined heuristically. For example, the α parameter, which controls the strictness of TE-dependence, may benefit from further optimization. Likewise, the correlation threshold (r = 0.5) used to identify task-induced ROI signals could be refined to improve specificity and sensitivity. Finally, the echo times (9.98, 21.61, and 33.24 ms) were selected based on commonly recommended values for BOLD sensitivity—particularly the optimal TE of approximately 30 ms—as well as constraints imposed by the MRI scanner. (Gowland, P. A., & Bowtell, 2007). While these values were expected to produce monotonically increasing BOLD signals across most gray matter voxels, anatomical variability may cause some voxels to deviate from this pattern. Future work will explore optimal TE selection tailored to individual brain structures to enhance BOLD component extraction.

## 5. Conclusion

In summary, our findings demonstrate that the BOLD-filter outperforms the typical preprocessing method (TYP) in analyzing task-based fMRI functional networks. The BOLD-filter effectively enhances sensitivity to task-induced brain activity, allowing for more reliable identification of activated regions and functional connectivity patterns. Notably, it revealed gender-related differences in processing everyday behaviors—differences that were not captured using TYP. These results suggest that the BOLD-filter has strong potential to uncover functional connectivity that may otherwise remain undetected, offering a promising tool for advancing the study of brain function through task-based fMRI.

### Author Contributions

Y-WS: Conceptualization, Data curation, Formal Analysis, Investigation, Methodology, Software, Project administration, Validation, Visualization, Writing – original draft, Writing –review & editing. U-SC: Conceptualization, Data curation, Investigation, Resources, Writing – original draft, Writing – review & editing. TT: Stimulation Design, Acquiring MRI data, Writing-review & editing. SO: Conceptualization, Data curation, Formal Analysis, Funding, Investigation, Project administration, Resources, Supervision, Writing – review & editing.

### Funding

This study was supported by JSPS KAKENHI Grant Number 21H02800, 21K11316 and 25K03427.

