## Supplementary Figures Methods for "Enhancing Functional Connectivity Analysis in Task-Based fMRI Using the BOLD-Filter Method: Greater Network and Activation Voxel Sensitivities"

### Supplementary Figure 1

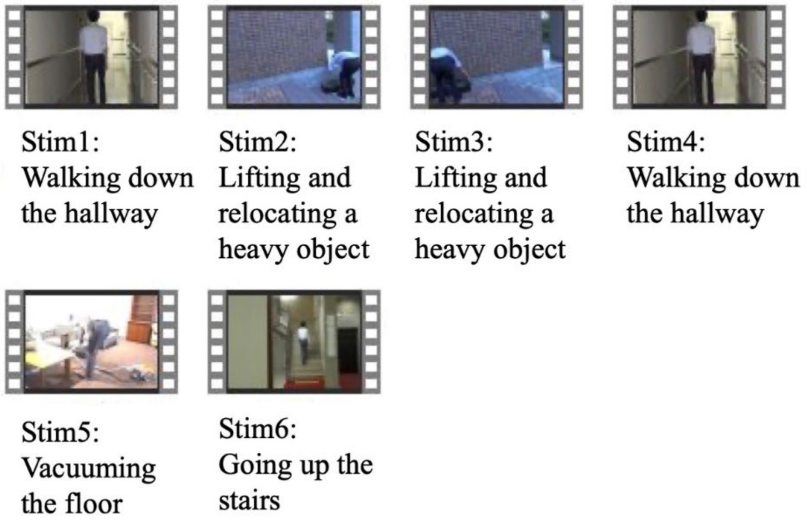


**Visual stimulation**

The video clips used for stimulation depicted everyday behaviors as follows: Stim1 and Stim4 represented "walking down the hallway," Stim2 represented "lifting and relocating a heavy object," Stim3 represented "going up the stairs," and Stim5 represented "vacuuming the floor."

### Supplementary Figure 2

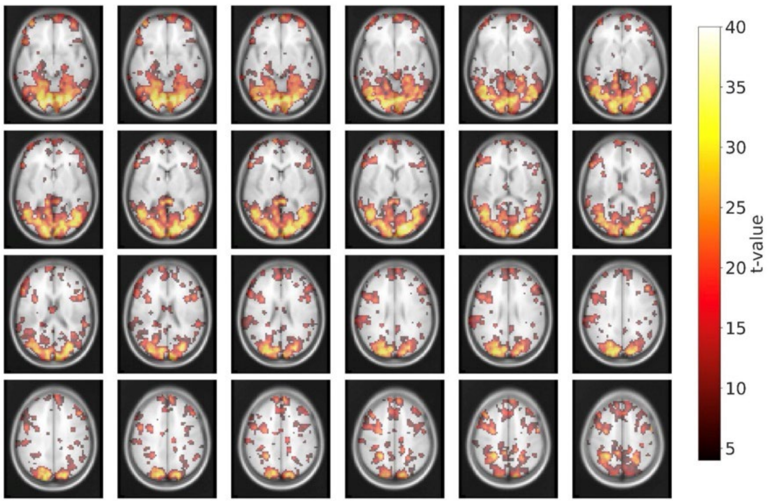


(a)

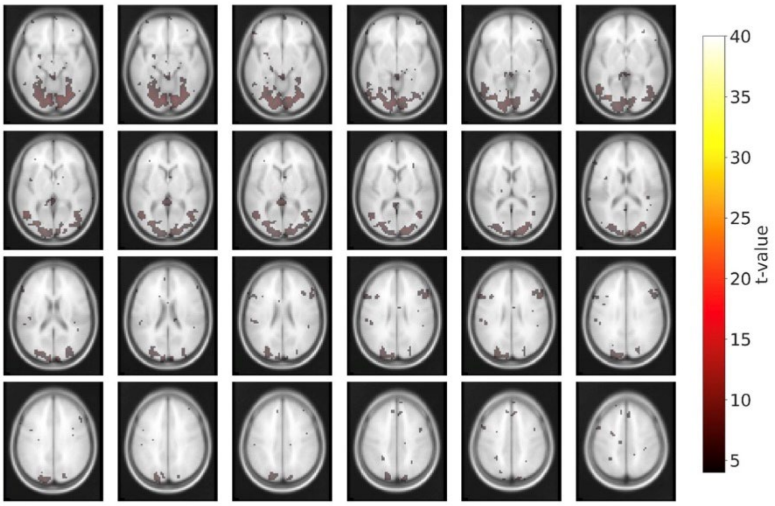


(b)

**Activation maps**

This figure presents activation maps from another subject at a low threshold (t = 4). The color bar indicates *t*-values. (a) Activation map obtained using BOLD-filter preprocessing (BOLDFLT); (b) Activation map obtained using conventional preprocessing (TYP).

**Supplementary Figure 3**

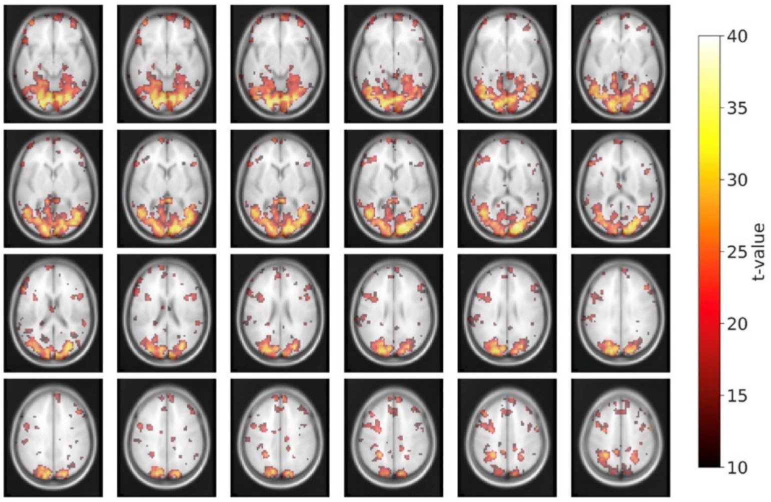


(a)

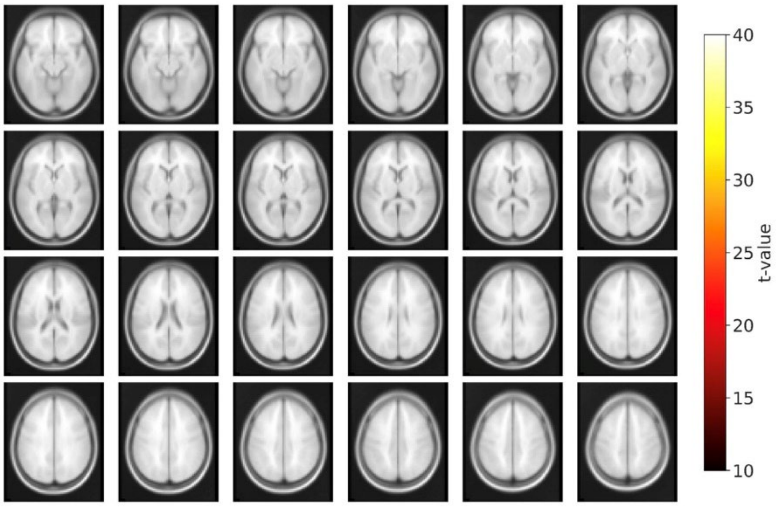


(b)

**Activation maps**

This figure presents activation maps from another subject at a high threshold (t = 10). The color bar indicates *t*-values. (a) Activation map obtained using BOLD-filter preprocessing (BOLDFLT); (b) Activation map obtained using conventional preprocessing (TYP).

**Supplementary Figure 4**

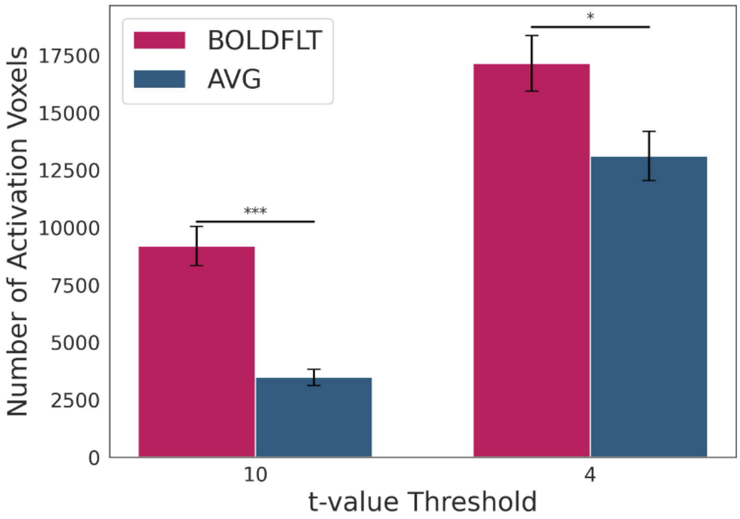


**Number of activation voxels**

Activation voxels were quantified for BOLD-filter preprocessing (BOLDFLT) and conventional preprocessing using the average of the three echo signals (AVG), at both high and low statistical thresholds. At the high threshold (t = 10), BOLDFLT produced approximately 2.6 times more activation voxels than AVG. At the low threshold (t = 4), BOLDFLT yielded over 1.3 times more activation voxels than AVG.

(a) BOLDFLT; (b) AVG.

**Supplementary Figure 5**

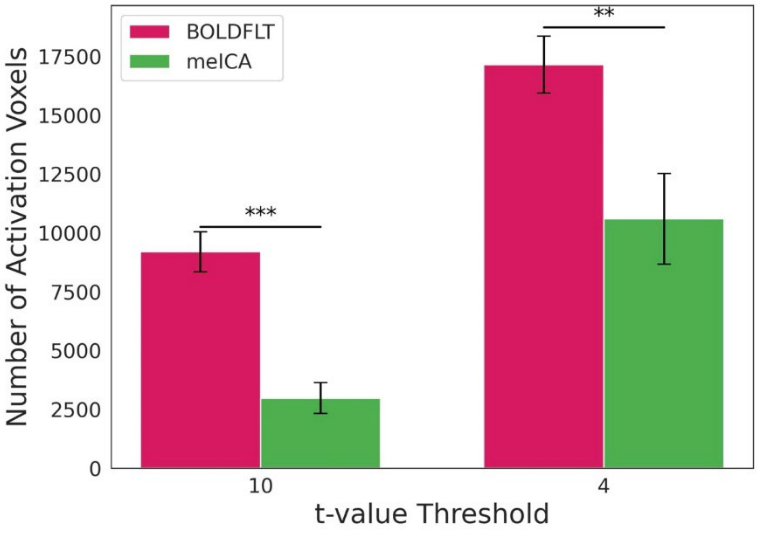


**Number of activation voxels**

Activation voxels were quantified for BOLD-filter preprocessing (BOLDFLT) and multi echo ICA preprocessing (meICA) of the three echo signals (meICA), at both high and low statistical thresholds. At the high threshold (t = 10), BOLDFLT produced approximately 3 times more activation voxels than meICA. At the low threshold (t = 4), BOLDFLT yielded over 1.6 times more activation voxels than meICA.

(a) BOLDFLT; (b) meICA.

**Supplementary Figure 6**

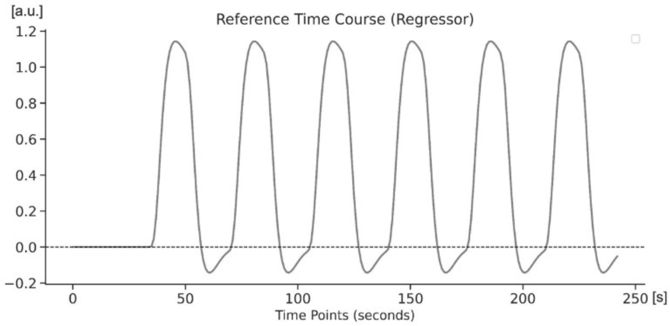


**Reference time course (i.e., Task regressor)**

The plot shows the regressor used in the general linear model (GLM), which was applied to the time series of each voxel. A two-gamma hemodynamic response function (HRF) was used to model the expected BOLD response to task stimulation.

**Supplementary Figure 7**

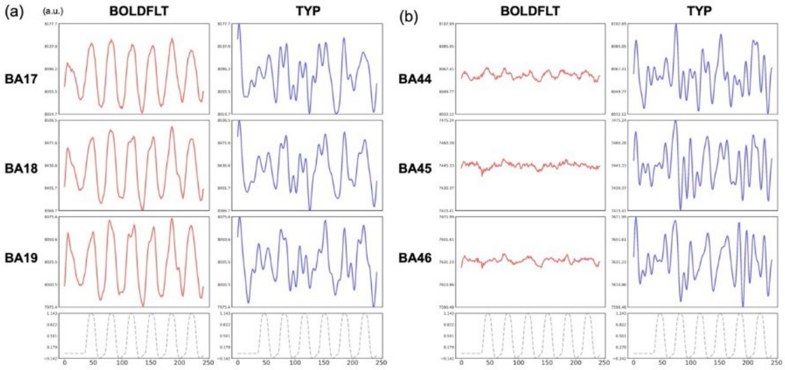


**Time course signals**

Time course signals were averaged across representative regions of interest (ROIs) in a single subject for both BOLD-filter preprocessing (BOLDFLT) and conventional preprocessing (TYP). Red lines represent BOLDFLT, blue lines represent TYP, and dashed lines indicate the reference time course (task regressor).

(a) Primary and secondary visual areas: Brodmann areas 17, 18, and 19

(b) Higher-order frontal regions: Brodmann areas 44, 45, and 46.

**Supplementary Figure 8**

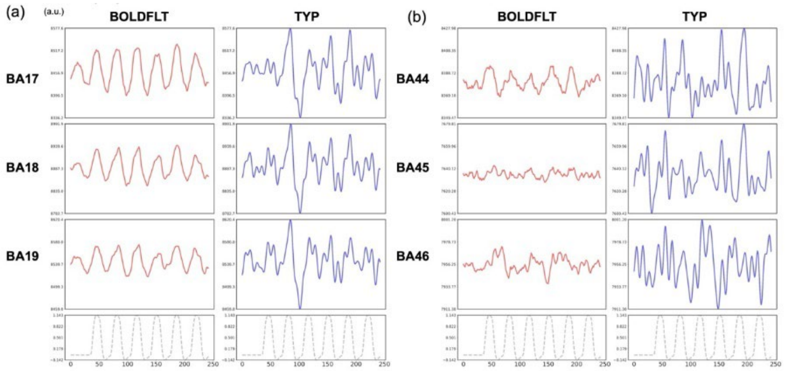


**Time course signals**

Time course signals were averaged across representative regions of interest (ROIs) in another single subject for both BOLD-filter preprocessing (BOLDFLT) and conventional preprocessing (TYP). Red lines represent BOLDFLT, blue lines represent TYP, and dashed lines indicate the reference time course (task regressor).

(a) Primary and secondary visual areas: Brodmann areas 17, 18, and 19

(b) Higher-order frontal regions: Brodmann areas 44, 45, and 46.

Supplementary Methods

```
%========================================
% Open-source MATLAB code for BOLD-filter
%========================================

%-------------------------------
% User-defined parameters
%-------------------------------
% 1. TE values (in ms)
TE1 = 9.98;
TE2 = 21.61;
TE3 = 33.24;

% 2. BOLD ratio threshold coefficient
alphaCoeff = 0.2;
alpha12 = 1 + alphaCoeff * (TE2 / TE1 - 1);
alpha23 = 1 + alphaCoeff * (TE3 / TE2 - 1);

% 3. T2* range thresholds (in ms)
T2_upper = 100;
T2_lower = 15;

% 4. Background intensity threshold (used for TE3)
thBackGround = 1000;

% 5. Input 4D data (row, col, slice, time) for each echo
dataTE1 = single(dataTE1);
dataTE2 = single(dataTE2);
dataTE3 = single(dataTE3);

%-------------------------------
% Preprocessing and Initialization
%-------------------------------
matrixSize = size(dataTE1);
[xRow, yColumn, Nslice, Nvol] = deal(matrixSize(1), matrixSize(2),
matrixSize(3), matrixSize(4));

% Ensure even number of volumes
if mod(Nvol, 2) == 1
    Nvol = Nvol - 1;
    dataTE1 = dataTE1(:, :, :, 1:Nvol);
    dataTE2 = dataTE2(:, :, :, 1:Nvol);
    dataTE3 = dataTE3(:, :, :, 1:Nvol);
end

% Initialize data containers
allDataSep = zeros(xRow, yColumn, Nslice, Nvol, 3, 'single'); % 3
echoes
buffOnes = ones(xRow, yColumn, Nslice, Nvol, 'single');
```

```
% Compute mean over time for each echo
avgDataTE1 = mean(dataTE1, 4);
avgDataTE2 = mean(dataTE2, 4);
avgDataTE3 = mean(dataTE3, 4);

% Populate averaged data
allDataSep(:,:,:,:,1) = buffOnes .* avgDataTE1;
allDataSep(:,:,:,:,2) = buffOnes .* avgDataTE2;
allDataSep(:,:,:,:,3) = buffOnes .* avgDataTE3;

% BOLD signal initialization
selDataBOLD = buffOnes .* avgDataTE3;

% Tracking maps
BOLDcntMap = zeros(xRow, yColumn, Nslice);
chkBOLDcntMap = zeros(xRow, yColumn, Nslice);

cntBack = 0;

%================================
% Main Processing Loop
%================================
for k = 1:xRow
    for l = 1:yColumn
        for m = 1:Nslice

            % Extract voxel time series for each TE
            testSig1 = squeeze(dataTE1(k,l,m,:));
            testSig2 = squeeze(dataTE2(k,l,m,:));
            testSig3 = squeeze(dataTE3(k,l,m,:));
            voxelMean = mean(testSig3);

            if voxelMean > thBackGround
                % Estimate R2* (in 1/s)
                R2 = 1000 * (log(mean(testSig1)) -
log(mean(testSig2))) / (TE2 - TE1);

                if R2 < (1000 / T2_lower) && R2 > (1000 / T2_upper)
                    cntBack = cntBack + 1;

                    % FFT transformation
                    y_testSig1 = fft(testSig1);
                    y_testSig2 = fft(testSig2);
                    y_testSig3 = fft(testSig3);

                    % Normalize magnitudes
                    y_DC1 = abs(y_testSig1(1));
                    y_DC2 = abs(y_testSig2(1));
                    y_DC3 = abs(y_testSig3(1));
```

```
                    % Frequency filtering
                    z_BOLD_pos = zeros(Nvol/2 - 1, 1);
                    for i = 2:Nvol/2 - 1
                        tmp_per1 = abs(y_testSig1(i)) / y_DC1;
                        tmp_per2 = abs(y_testSig2(i)) / y_DC2;
                        tmp_per3 = abs(y_testSig3(i)) / y_DC3;

                        ratio12 = tmp_per2 / tmp_per1;
                        ratio23 = tmp_per3 / tmp_per2;

                        % Frequency selection criteria
                        cond1 = (ratio12 > alpha12) && (ratio23 >
alpha23);
                        sameRealSign = all(sign(real([y_testSig1(i),
y_testSig2(i), y_testSig3(i)])) == sign(real(y_testSig1(i))));
                        sameImagSign = all(sign(imag([y_testSig1(i),
y_testSig2(i), y_testSig3(i)])) == sign(imag(y_testSig1(i))));
                        cond3 = sameRealSign && sameImagSign;

                        if cond1 && cond3
                            z_BOLD_pos(i-1) = 1;
                        end
                    end

                    % Mirror frequency for negative band
                    z_BOLD_neg = flip(z_BOLD_pos);
                    tmp_CutFrq = [0; z_BOLD_pos; 0; z_BOLD_neg];

                    % Apply frequency mask
                    y_bold_sig1 = y_testSig1 .* tmp_CutFrq;
                    y_bold_sig2 = y_testSig2 .* tmp_CutFrq;
                    y_bold_sig3 = y_testSig3 .* tmp_CutFrq;

                    % Restore DC components
                    y_bold_sig1([1, Nvol]) = y_testSig1([1, Nvol]);
                    y_bold_sig2([1, Nvol]) = y_testSig2([1, Nvol]);
                    y_bold_sig3([1, Nvol]) = y_testSig3([1, Nvol]);

                    % Inverse FFT to obtain filtered signal
                    z_bold_sig1 = ifft(y_bold_sig1);
                    z_bold_sig2 = ifft(y_bold_sig2);
                    z_bold_sig3 = ifft(y_bold_sig3);

                    % Artifact reduction using correlation
                    fixSig1 = abs(z_bold_sig1);
                    fixSig2 = abs(z_bold_sig2);
                    fixSig3 = abs(z_bold_sig3);

                    corr12 = abs(corr(fixSig1, fixSig2));
```

```
                    corr23 = abs(corr(fixSig2, fixSig3));

                    % Frequency count and filtering conditions
                    freqCount = sum(tmp_CutFrq > 0);
                    BOLDcntMap(k,l,m) = freqCount;
                    chkBOLDcntMap(k,l,m) = freqCount;

                    fixCond1 = freqCount > 2;
                    fixCond2 = (corr12 < 0.995) && (corr23 < 0.995);
                    fixCond3 = (corr12 > 0.5) && (corr23 > 0.5);

                    if fixCond1 && fixCond2 && fixCond3
                        selDataBOLD(k,l,m,:) = abs(z_bold_sig3);
                        allDataSep(k,l,m,:,1) = z_bold_sig1;
                        allDataSep(k,l,m,:,2) = z_bold_sig2;
                        allDataSep(k,l,m,:,3) = z_bold_sig3;
                    else
                        selDataBOLD(k,l,m,:) = DC3;
                        allDataSep(k,l,m,:,1) = DC1;
                        allDataSep(k,l,m,:,2) = DC2;
                        allDataSep(k,l,m,:,3) = DC3;
                        chkBOLDcntMap(k,l,m) = 0;
                    end
                end
            end
        end
    end
end

% Final output
allDataSep = abs(allDataSep); % Convert complex to magnitude
BOLDsig = allDataSep(:,:,:,:,3); % Extracted BOLD component

% A high-pass filter with a cutoff frequency of 0.008 Hz (using
spm_filter)
% was applied to the final data 'BOLDsig' prior to further
processing.
```
